# Paternal chromosome elimination and X non-disjunction on asymmetric spindles in *Sciara* male meiosis

**DOI:** 10.1101/2021.05.13.444088

**Authors:** Brigitte de Saint Phalle, Rudolf Oldenbourg, Donna F. Kubai, E. D. Salmon, Susan A. Gerbi

## Abstract

Meiosis in male *Sciara* is unique with a single centrosome. A monopolar spindle forms in meiosis I, but a bipolar spindle forms in meiosis II. The imprinted paternal chromosomes are eliminated in meiosis I; there is non-disjunction of the X in meiosis II. Despite differences in spindle construction and chromosome behavior, both meiotic divisions are asymmetric, producing a cell and a small bud. Observations of live spermatocytes made with the LC-PolScope, differential interference contrast optics and fluorescence revealed maternal and paternal chromosome sets on the monopolar spindle in meiosis I and formation of an asymmetric monastral bipolar spindle in meiosis II where all chromosomes except the X congress to the metaphase plate. The X remains near the centrosome after meiosis I and stays with it as the spindle forms in meiosis II. Electron microscopy revealed amorphous material between the X and the centrosome. Immunofluorescence with an antibody against the checkpoint protein Mad2 stains the centromeres of the maternal X dyad in late meiosis I and in meiosis II where it fails to congress to the metaphase plate. Mad2 is also present throughout the paternal chromosomes destined for elimination in meiosis I, suggesting a possible role in chromosome imprinting. If Mad2 on the X dyad mediates a spindle checkpoint in meiosis II, it may delay metaphase to facilitate formation of the second half spindle through a non-centrosomal mechanism.

## Introduction

The precision of chromosome separation has fascinated biologists for over a century. Typically, the duplicated centrosomes move apart from one another to each end of the bipolar spindle which forms. The chromosomes then segregate equally to the two spindle poles. Much can be learned about the underlying mechanism by perturbing the normal process. This has been done by others by chemical or genetic means. The unique meiotic divisions in spermatocytes of the lower dipteran fly *Sciara* offer a naturally occurring variant for chromosome division and is the subject of this paper.

Each spermatocyte produces a single sperm in *Sciara* instead of the usual four. This sperm contains the maternal autosomes, both copies of the maternal X chromosome and germ-line limited “L” chromosomes (Metz, 1938; Gerbi, 1986; Goday and Esteban, 2001). How this occurs through meiosis is diagrammed in Figure 1. There is no synapsis of the maternal and paternal homologues in prophase of male meiosis I (Metz et al., 1926). Spermatocytes enter meiosis with a single centrosome, which produces a monopolar spindle (Figure 1A). No metaphase plate is formed. The chromosomes are imprinted (a term first coined in *Sciara*; Crouse, 1960) with regard to their parental origin. All the maternal chromosomes as well as the L chromosomes (which escape the parental imprint; Crouse et al., 1971) are seen near the single centrosome. The paternal chromosomes are distal to the centrosome and are excluded in a bud of cytoplasm (Figure 1B).

**Figure 1.**
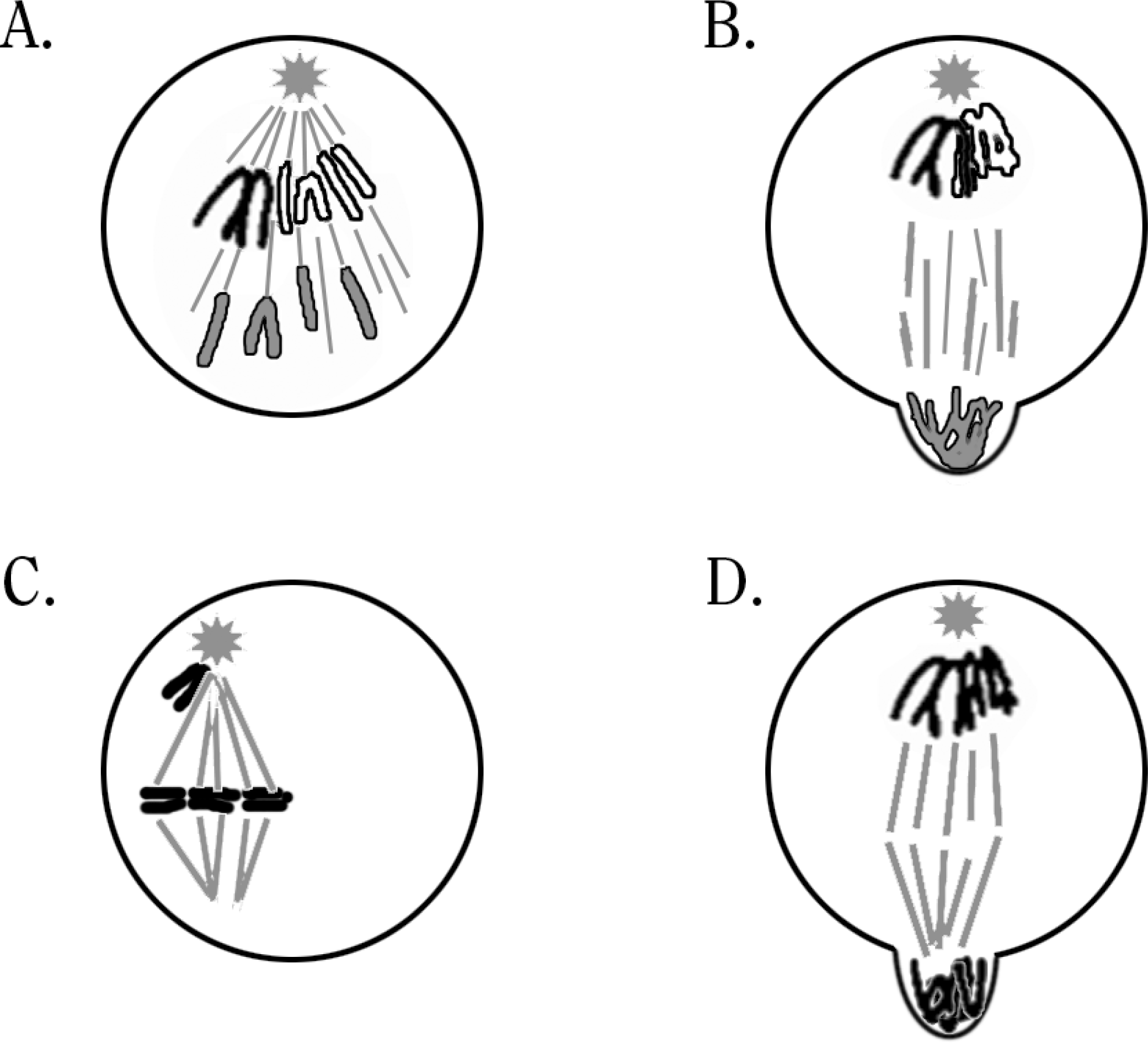
Cytology of Sciara male meiosis I and meiosis II (modified after DuBois, 1933). **A-B:** Meiosis I. Maternal chromosomes white outlined in black, paternal chromosomes grey outlined in black, L’s black, microtubules and centrosome aster grey. Grey star represents single centrosome. **A.** Anaphase-like separation of the maternal and paternal homologues on the monopolar spindle without congression to a metaphase plate. The L chromosomes go to the pole with the chromosomes of maternal origin. **B.** Telophase-like formation of two division products with a central spindle of microtubules separating them. The division product without a centrosome is budding out of the cell. **C-D:** Meiosis II. Chromosomes black, microtubules grey. **C.** Metaphase with X dyad at the centrosome, all other chromosomes at the metaphase plate. **D.** Telophase with the large central spindle separating the division products. The nullo-X division product, which lacks a centrosome, is budding out of the cell.

In Meiosis II (Figure 1 C-D) the chromosomes congress to the metaphase plate of a bipolar spindle except for the X dyad that is seen prematurely at one pole (Figure 1C). The subsequent cell division is asymmetric (Figure 1D), and the nullo-X product of the division is cast off in a small bud (Metz, 1938).

The single cell that results from spermatogenesis has two copies of the X chromosome. Therefore, fertilization of the haploid oocyte by sperm containing two X chromosomes results in a zygote with three Xs. The cell corrects this problem by elimination of one or two X chromosomes in early cleavage divisions, depending on the sex of the embryo (Metz, 1938; Gerbi, 1986; Goday and Esteban, 2001; de Saint Phalle and Sullivan, 1996). The “controlling element” proximal to the X determines its behavior during male meiosis II (non-disjunction) and in early embryogenesis (chromosome elimination) (Crouse, 1960). The L chromosomes are also eliminated from the somatic lineage in embryogenesis.

The unique meiotic divisions in male *Sciara* raise several questions. Why is a monopolar spindle formed in meiosis I and a bipolar spindle in meiosis II, even though both spindles have only a single centrosome? How are the chromosomes imprinted? How do the chromosomes move on the monopolar spindle? What causes non-disjunction of the X dyad in meiosis II?

We have taken advantage of new technology in polarization microscopy, the liquid crystal (LC)-PolScope (Oldenbourg and Mei, 1995), which provided an opportunity to observe both meiotic divisions in living spermatocytes from *Sciara*. These observations were coupled with differential interference contrast (DIC) and fluorescence microscopy of living spermatocytes. The data were expanded by electron microscopy and immunofluorescence of fixed spermatocytes to begin to elucidate the underlying mechanisms for these unique meiotic divisions. Our data suggest that the chromosomes do not move on the monopolar spindle of meiosis I. The X dyad does not align on the metaphase plate nor does it move in meiosis II; instead, it is juxtaposed to the single centrosome and appears to be carried with it as the centrosomal half-spindle elongates. Subsequently, a second but shorter half-spindle forms on the other side of the metaphase plate, apparently by a non-centrosomal mechanism. The checkpoint protein Mad2 is located on the X centromere at the end of meiosis I and in meiosis II where it may delay metaphase sufficiently to allow the second half-spindle to form. Intriguingly, Mad2 also localizes to the arms of the paternal chromosomes in the anaphase-like stage of meiosis I, suggesting it may play a second role related to chromosome imprinting.

## RESULTS

### Meiosis I and II in Living Spermatocytes

Observations made on living cells from testes dissected into hemolymph or insect Ringer’s buffer were comparable. There is fairly good synchrony between the cells of one cyst but not between cysts of the same testis (Phillips, 1970). Cytoplasmic bridges allow communication between the cells within one cyst. The cells are all oriented with the buds being eliminated into the lumen of the cyst (Metz et al., 1926; Amabis et al., 1979; Kubai, 1982).

### Prophase I and prophase II

In prophase of meiosis I (PI), the nucleus contains the unpaired maternal and paternal X chromosomes and autosomes plus two L chromosomes. The chromosomes are not tightly condensed except the L chromosomes (Figure S1 A-C). In prophase of meiosis II (PII) the chromosomes are condensed more tightly than in PI and the L chromosomes are no longer differentially condensed compared to the other chromosomes (Figure S1 D-F). The cell size is about the same in II and PII, but the nucleus occupies 59% of the cell in PI yet only 21% in PII (Table S1), reflecting in part the fewer number of chromosomes.

### Meiosis I

The spindle microtubules could be visualized with polarization optics of living *Sciara* spermatocytes without staining, but the chromosomes, particularly in meiosis I, were often obscured by the highly birefringent mitochondria (Figure 2).

**Figure 2.**
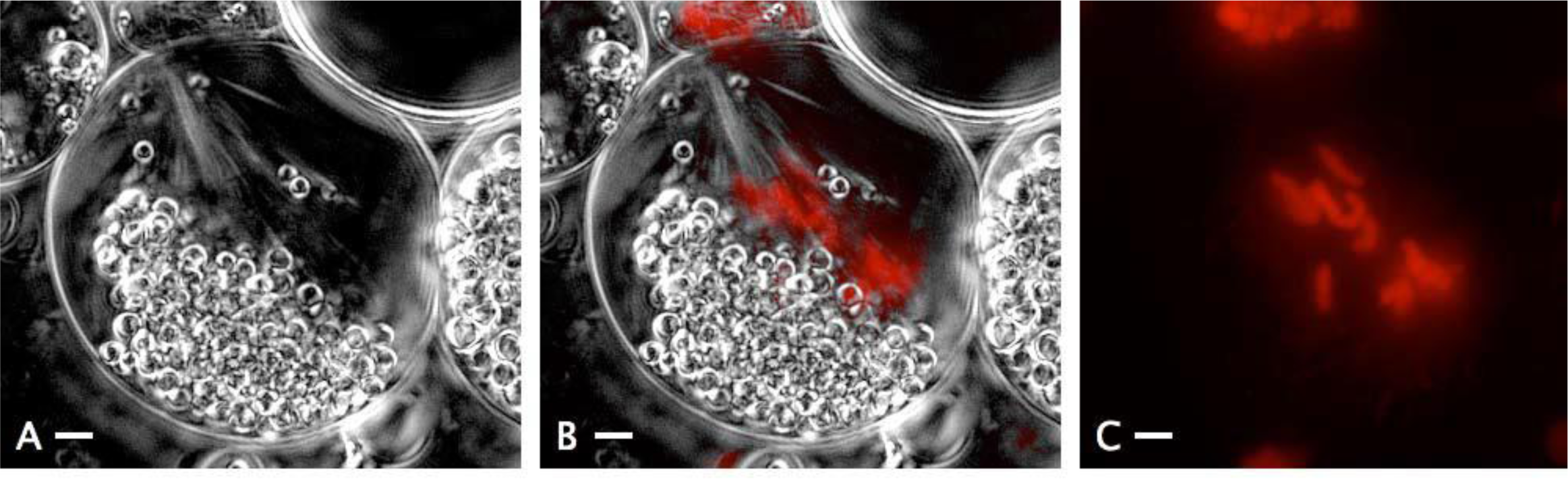
Meiosis I. Live LC-PolScope images and epifluorescence images of DNA stained with Hoechst. In meiosis I the paternal and maternal homologues are seen separating on a monopolar spindle. The maternals and Ls are above and to the left. All bars = 2 μm. **A.** LC-PolScope image is a projection of two separate slices of a POL stack chosen to show some of the monopolar spindle while reducing the area of mitochondria which usually completely obscure the chromosomes. **B.** LC-PolScope image combined with Hoechst image of DNA (red). **C.** Image of DNA stained with Hoechst is a projection of a full stack of DNA images.

We used the vital stain Hoechst 33342 to stain the DNA so we could image chromosomes through the mitochondria by epifluorescence.

In meiosis I the spindle is monopolar and the chromosomes do not congress to a metaphase plate. Instead, the cells in meiosis I move directly from prophase (Figure S1 A-C) to an anaphase configuration where the maternal homologues and L chromosomes are seen near the single centrosome and the paternal homologues are at the opposite end of the cell (Figure 2). Despite our ability to visualize chromosome movement in meiosis II, we never observed significant chromosome movement in the anaphase-like state of meiosis I, thus calling into question whether chromosomes move on the monopolar spindle.

### Meiosis II

Polarization microscopy of spermatocytes in prometaphase of meiosis II reveals birefringence suggestive of a monopolar spindle nucleated by the single centrosome (Figure 3 B-E). This half-spindle lengthens over time, and by metaphase II a second but shorter half-spindle forms on the other side of the metaphase plate to create an asymmetric bipolar (but monocentric) spindle (Figure 4). The X dyad does not congress to the metaphase plate like the other chromosomes. Instead, the X dyad is found at the centrosomal pole and moves with it as the half-spindle elongates in prometaphase II (Figure 3). Thus, by DIC microscopy, the distance between the X dyad and the metaphase plate increases during the progression of prometaphase II (Figure 3A and 3F) though the distance between the X dyad and the single centrosome remains unchanged as viewed with polarization optics (Figure 3 B-E). Only a few exceptions were seen with the LC-PolScope where an X could not be seen at the centrosomal pole or in one case where the X was seen stretched between the metaphase plate and the centrosomal pole (data not shown).

**Figure 3.**
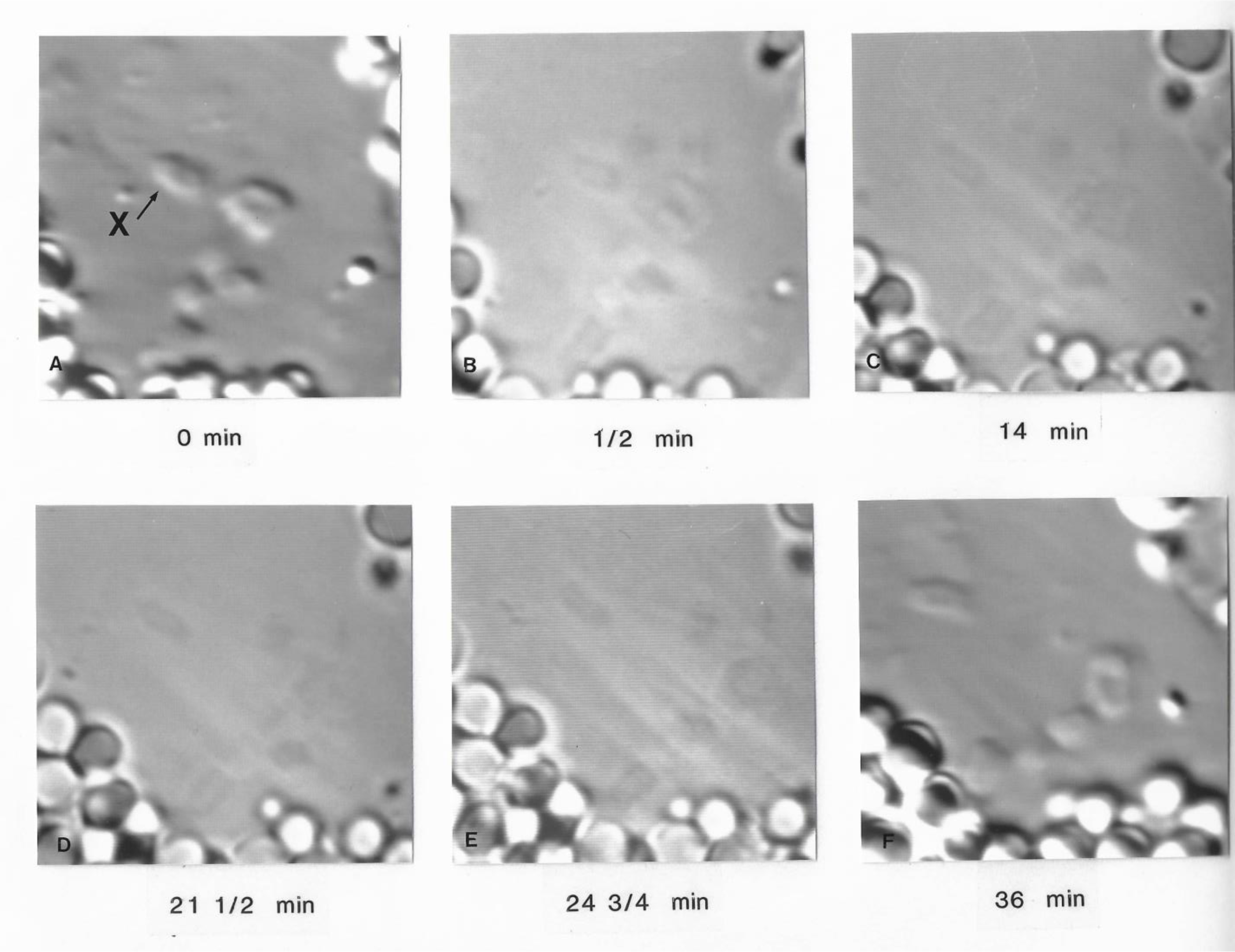
Location of the X dyad in meiosis II. A spermatocyte viewed with DIC optics (panels A and F) and standard polarization optics (panels B-E) while it enters meiosis II: prometaphase (A-B) and metaphase (C-F). **A.** 0 min. **B.** 0.5 min. **C.** 14 min. **D.** 21.5 min. **E.** 24.75 min. **F.** 36 min.

**Figure 4.**
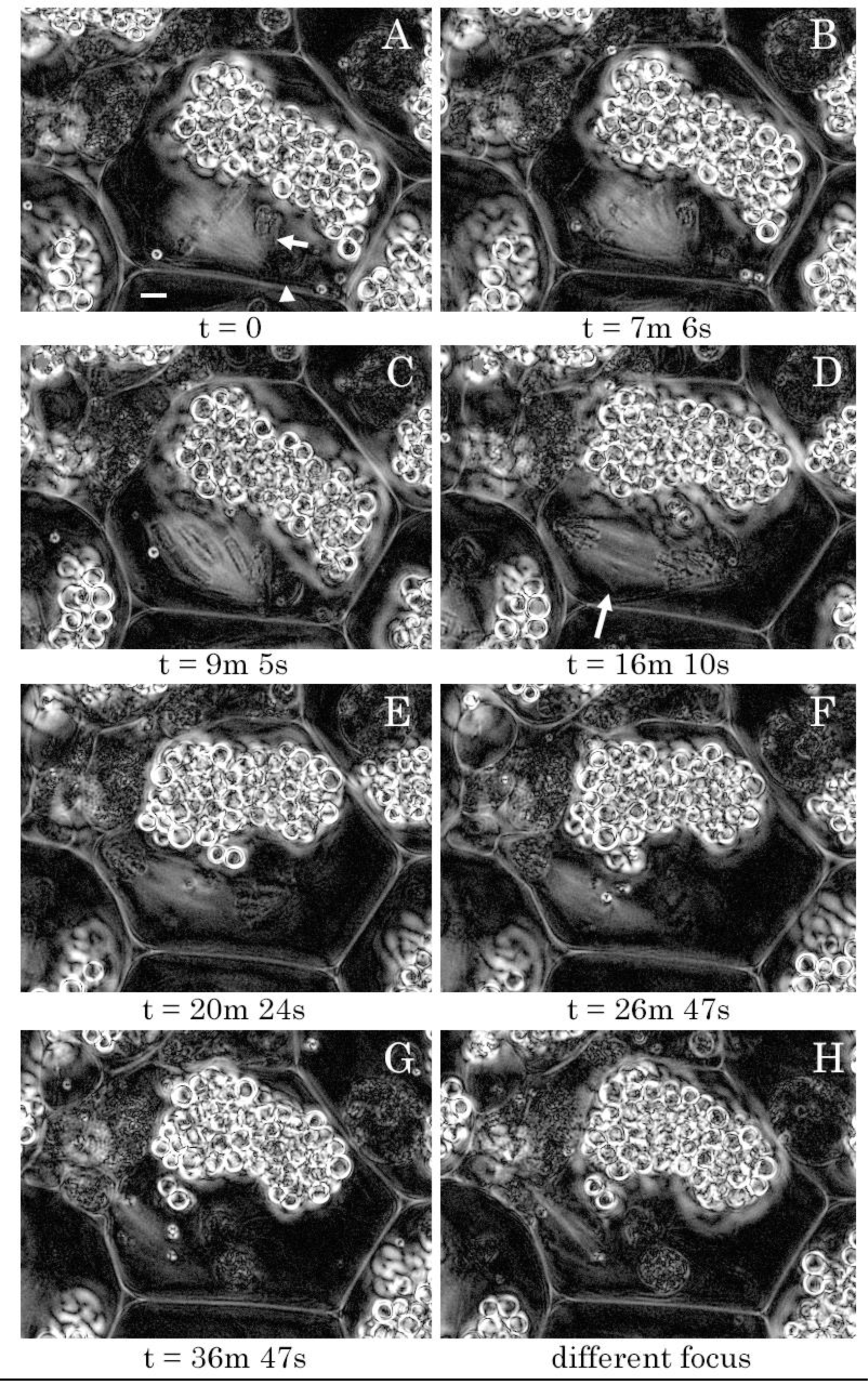
Meiosis II metaphase through telophase with budding attempt. **A.** t = 0. Metaphase with the chromosomes aligned on a conventional metaphase plate, centrosome location marked by arrowhead (the centrosome is not in the plane of focus). The X dyad (arrow) does not congress to the metaphase plate and is near the centrosome. **B.** t = 7m 6s. Start of anaphase. **C.** t = 9m 5s. Mid-anaphase. **D.** t = 16m 10s. Late anaphase, an arrow marks the central spindle. **E.** 20m 24s. The division product without a centrosome has contacted the cell membrane and is making a bulge in it. **F.** t = 26m 47s. The budding division product is in a pocket of cell membrane (arrow). **G.** 36m 47s. The bud is forming. **H.** In a different focal plane than panel G, a bundle of microtubules can be seen extending into the bud neck. During meiosis II the nucleus with a centrosome never contacted the cell membrane.

Anaphase II on this asymmetric bipolar spindle is conventional (Figure 4 A-D). By telophase II, the chromosomes including the X dyad are tightly clustered at the single centrosomal end of the spindle, though the chromosomes destined to be eliminated are not as tightly clustered (Figure 4 E-H). On several occasions we saw the division product with no centrosome contact the cell membrane and indent the cell membrane, whereas the division product with the centrosome never contacts the cell membrane (Figure 4 E-H). This is the beginning of bud formation where the nullo-X group of chromosomes are eliminated in the bud. By late anaphase and telophase of meiosis II, the half-spindle closest to the single centrosome dissipates, whereas the half-spindle distal to the single centrosome elongates and becomes more birefringent. Figure 4G shows some of the spindle microtubules extending through the bud neck, which is better visualized in a different focal plane (Figure 4H).

### Electron Microscopy of Meiosis II

Electron microscopy of serial sections through spermatocytes in meiosis II revealed kinetochores on the chromosomes on the metaphase plate but not on the X dyad that fails to align on the metaphase plate (Figure 5). It is known from c-banding that the centromere end of the X dyad is distal to the metaphase plate (Abbott and Gerbi, 1981). Although a distinct kinetochore fails to form at the X centromere, amorphous material is seen beyond the X centromere region and the centrosome (Figure 5).

**Figure 5.**
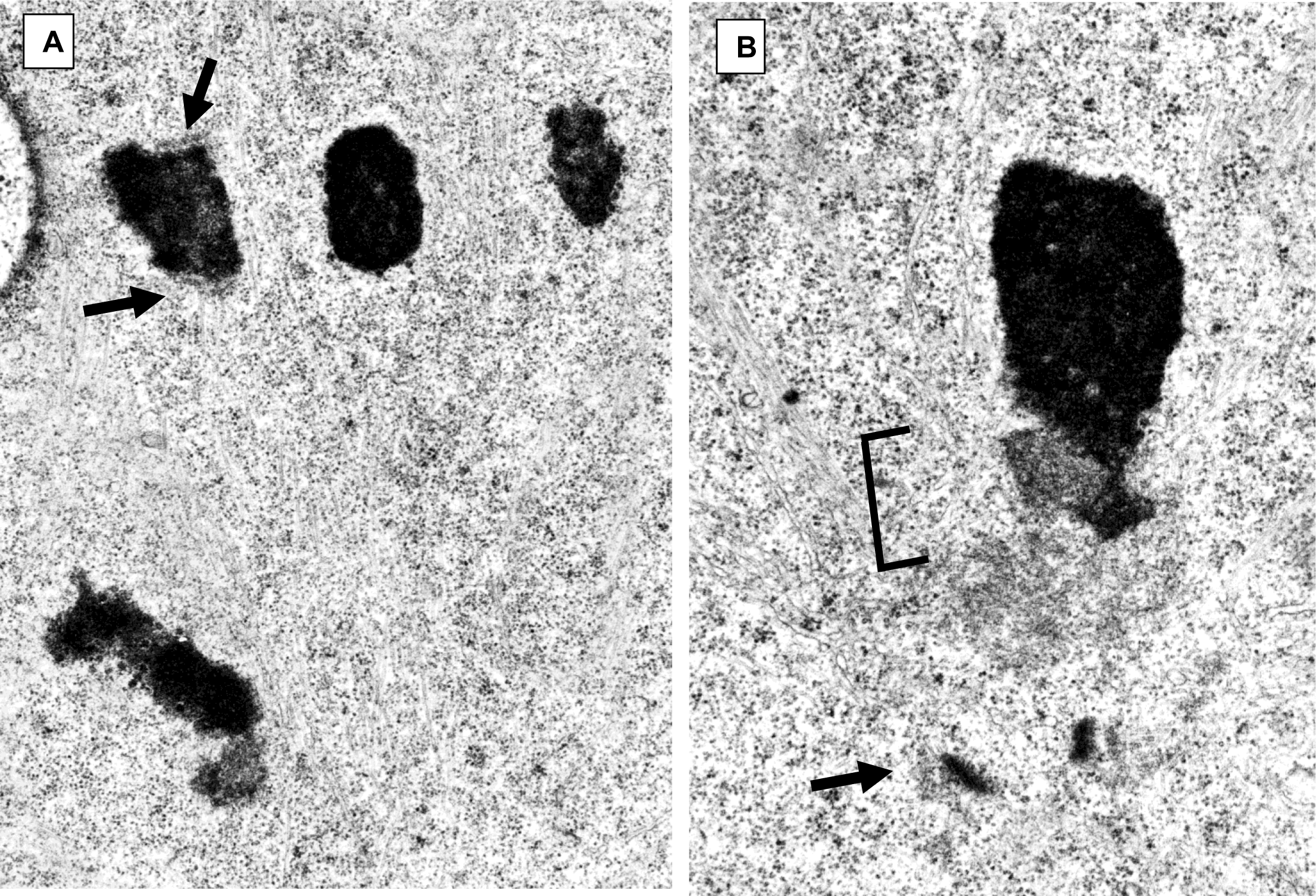
Ultrastructure of chromosomes in meiosis II. A. Three chromosomes are on the metaphase plate with well-formed kinetochores (arrows) seen on one of them in this section, and microtubules emanating from the kinetochores. No canonical kinetochore was ever seen on the X dyad. **B.** The X dyad (X) is found close to the centrosome (arrow) with amorphous material in-between (bracket).

### Localization of Mad2 in Sciara male meiosis

Mad2 associates with kinetochores until they capture microtubules (Waters et al., 1999). It serves as a checkpoint to delay progression into anaphase until all chromosomes are tethered onto microtubules. We carried out immunofluorescent staining with the R793 polyclonal antibody against *Drosophila* Mad2 to investigate the localization of this check-point protein in *Sciara* spermatocytes since they lack metaphase in meiosis I and the X dyad does not align on the metaphase plate in meiosis II.

### Mad2 staining of meiosis I spermatocytes

In the anaphase-like configuration of the monopolar spindle in meiosis I, the maternal and L chromosomes that are near the single centrosome do not exhibit Mad2 staining but the paternal chromosomes have Mad 2 associated with chromosome arms (Figure 6 A-C). Even though this localization is not appropriate for the checkpoint function which would involve the kinetochore, it indicates a difference in protein associations between the paternal chromosomes and the other (maternal + L) chromosomes. This may reflect their differential parental imprint.

**Figure 6.**
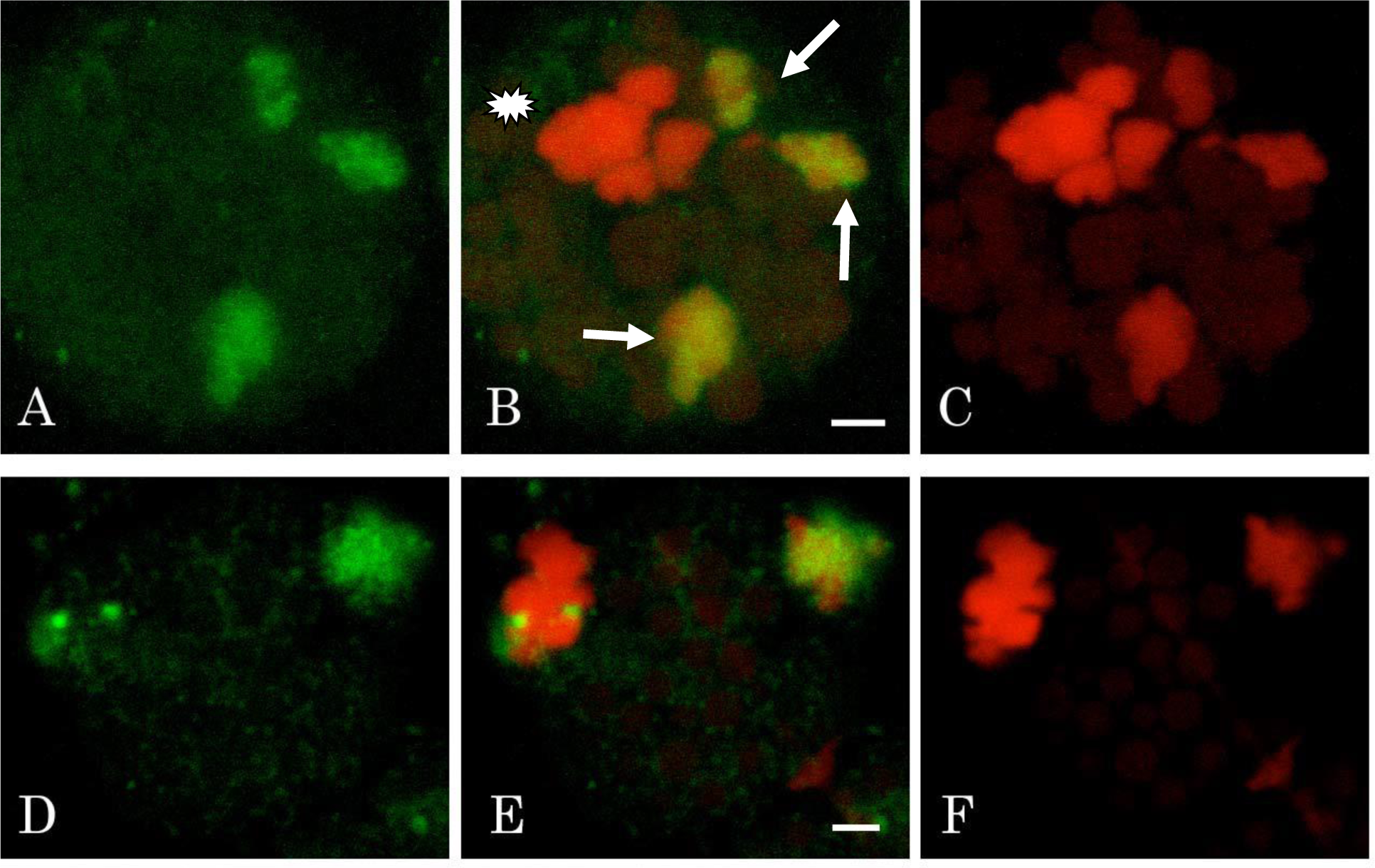
Mad2 staining in meiosis I anaphase-like and telophase-like stages. Mad2 green, DNA (propidium iodide) red, bars = 2 μm. **A-C:** Maximum projection of 11 slices of a Z-stack of an anaphase-like spermatocyte in meiosis I. The maternal and L chromosomes, identifiable by their location near the centrosome (white asterisk), do not exhibit Mad2 staining but the fanned out paternal chromosomes (white arrows) have Mad 2 associated with chromosome arms. **D-F:** Telophase-like spermatocyte in meiosis I. The maternal division product associated with the centrosome has Mad2 dots and the paternal division product has Mad2 loosely associated with it.

In the telophase-like stage of meiosis I, there is a cloud of Mad2 staining over the paternal chromosomes to be discarded in a bud (Figure 6 D-F). In addition, two adjacent dots of Mad2 staining are seen in the cluster of maternal and L chromosomes next to the centrosome (Figure 6 D-F). Based on data from meiosis II (below), these two dots of Mad2 may be on the centromeres of the X dyad. Mad2 would not normally be present in telophase of mitosis (it dissociates from the kinetochore after microtubule capture (Waters et al., 1999) but has been less studied in meiosis).

### Mad2 stains the centrosome and X centromere in meiosis II

In spermatocytes near metaphase II, Mad2 marks the centromere of the X dyad near a larger, more diffuse area of Mad2 staining, which is probably the centrosome (Figure 7). This X dyad centromeric staining is reminiscent of the two dots of staining over the maternal set of chromosomes in the telophase-like stage of meiosis I.

**Figure 7.**
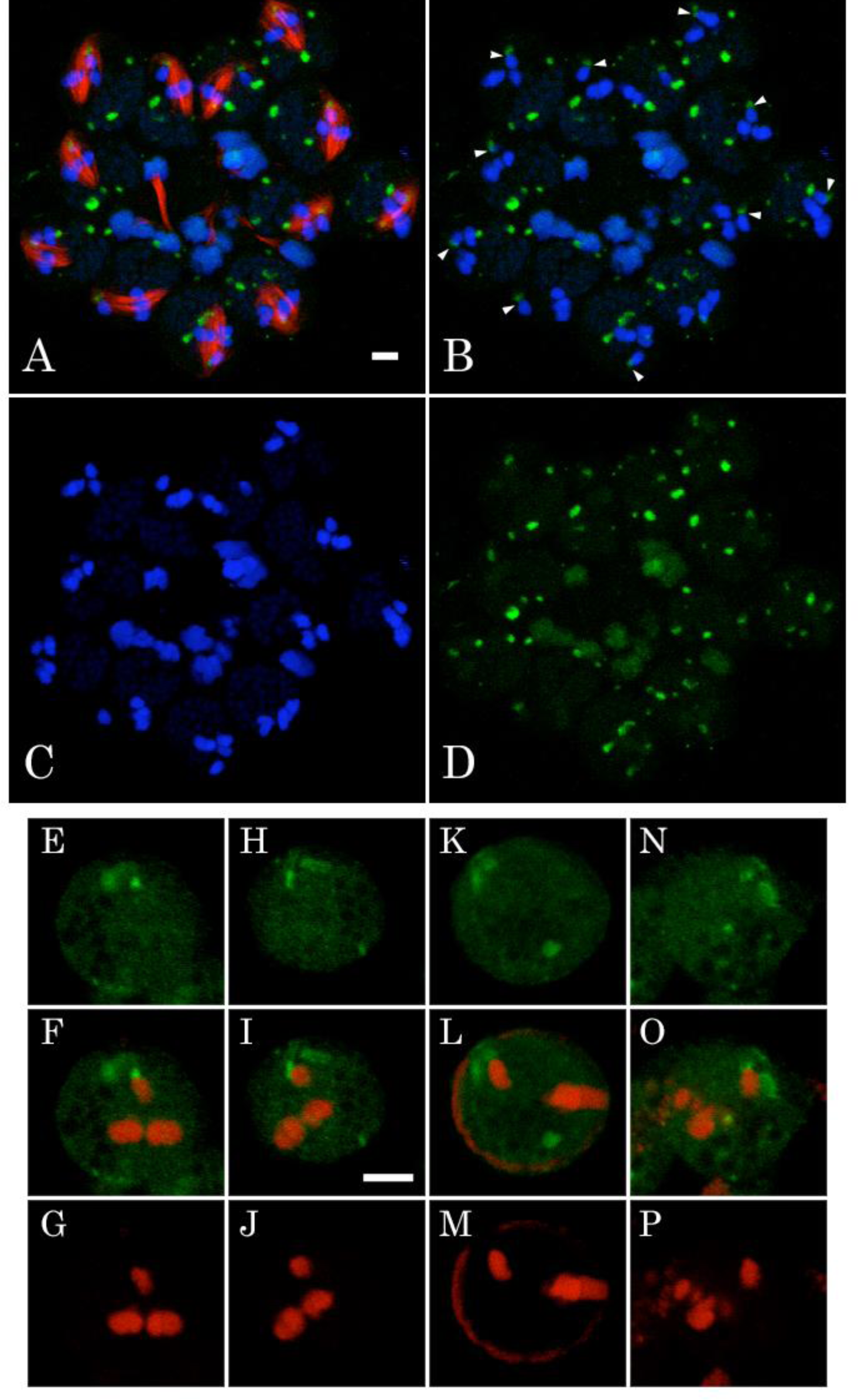
Mad2 staining in metaphase II. All bars = 2 μm. **A-D:** Maximum projection of 5 slices of a Z-stack of a group of spermatocytes in metaphase II, Mad2 green, DNA (Hoechst) blue, tubulin red. Mad2 stains the centromeres (arrowheads) of the X dyad, as well as nuclear foci not associated with chromosomes, the centrosome or spindle pole distal to the centrosome. The projection of 5 slices displays many of these Mad2 foci. **E-P:** Single spermatocytes near metaphase II. Mad2 green, DNA (propidium iodide) red. Bar = 2 μm, applies to all images. Double stain shows Mad2 marking the centromere of the X dyad near a larger, more diffuse area of Mad2 staining, probably the centrosome.

Figure 8 shows anaphase and telophase of meiosis II. In early anaphase, Mad2 persists on the X dyad (Figure 8 A-D). By late anaphase II (Figure 8 E-H) and telophase II (Figure 8 I-L), Mad2 staining is no longer seen over the X dyad but extraneous Mad2 foci are prominent. By late telophase II, the half spindle ending with the double-X set of chromosomes dissipates, but the spindle microtubules of the other half-spindle elongate into the bud with the nullo-X nucleus.

**Figure 8.**
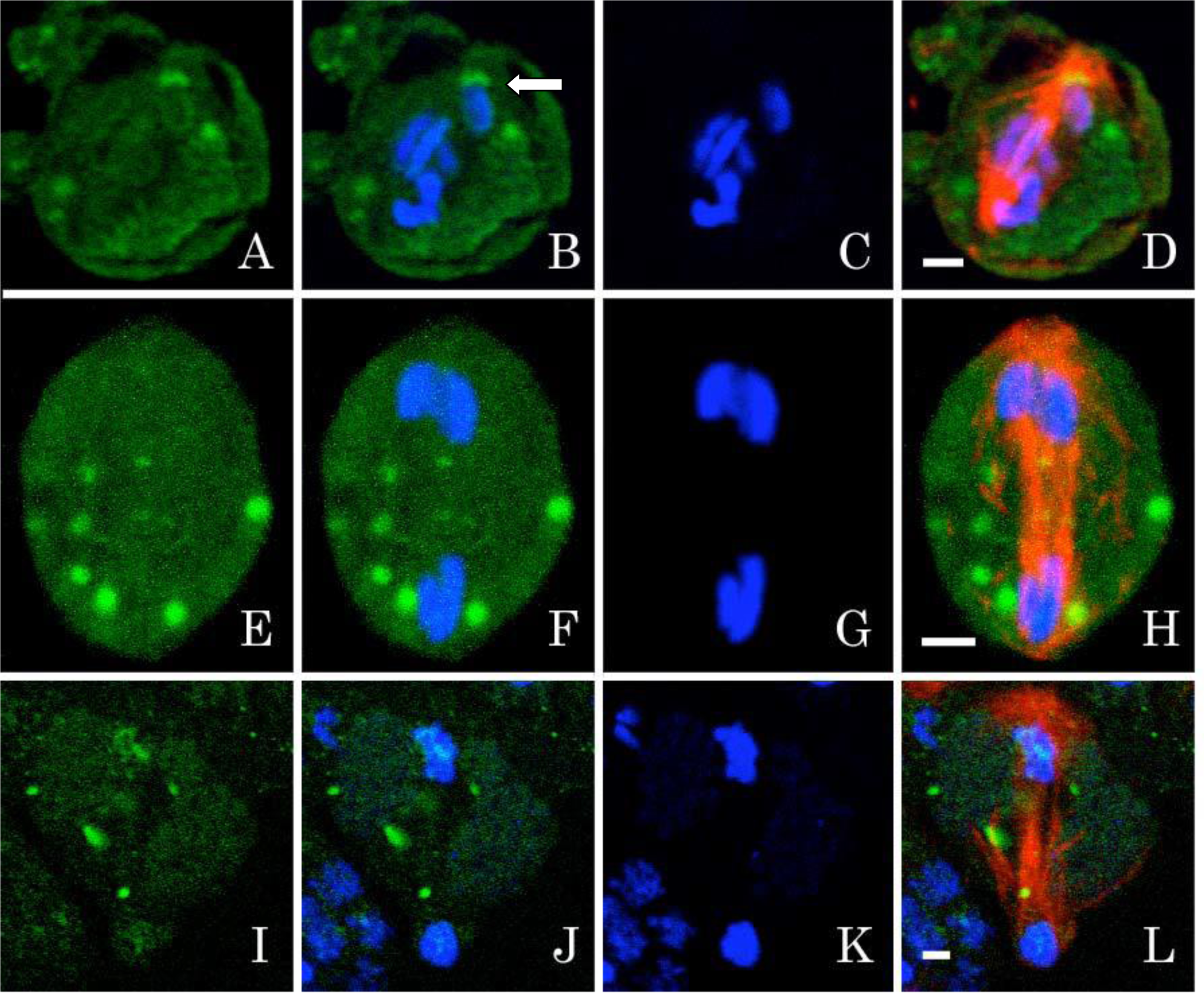
Mad2 staining in anaphase II and telophase II. Mad2 green, DNA (Hoechst) blue, tubulin red, all bars = 2 μm. **A-D:** Early anaphase II with Mad2 (white arrow) still on the X dyad and extraneous Mad2 foci. **E-H:** Maximum projection of 13 slices of a Z-stack so that all Mad2 foci are displayed. Late anaphase with the nullo X division product at the cell membrane. Mad2 no longer marks the X dyad; some extraneous Mad2 foci are prominent. **I-L:** Spermatocyte in telophase II with extraneous Mad2 foci. The division product without the X dyad is in a pouch of the cell membrane that will be budded off. The central spindle extends into the forming bud.

## DISCUSSION

*Sciara* executes its non-Mendelian meiosis with military precision: there is a monopolar spindle, the paternal chromosomes are eliminated and there is a non-disjunction of X in meiosis II. Elucidation of the underlying mechanisms for this programmed alternate version of a canonical meiosis may expand our understanding of universal principles.

### Asymmetric Spindles in Male *Sciara* Meiosis

The monopolar spindle in meiosis I and asymmetric spindle in meiosis II of *Sciara* spermatocytes both reflect the occurrence of only a single centrosome. The single centrosome is derived from giant centrioles in *Sciara’s* spermatogonial cells (Phillips, 1966 and 1967). Electron microscopy of early prophase spermatocytes from the related sciarid, *Trichosia,* showed that the two centrioles separate but are not associated with daughter centrioles; one centriole forms an aster and becomes the single spindle pole while the other centriole degenerates (Fuge, 1994). Typically, the centrosome is the major microtubule organizing center from which the spindle microtubules emanate (Mitchison and Kirschner, 1984). Therefore, a monopolar spindle would be expected to arise when there is only a single centrosome, which can be artificially induced in other systems by mutation, depletion or inhibition of components needed for centrosome duplication and separation (Wang et al 2014). Classic examples include the drug monastrol that inhibits the kinesin Eg5 needed for centrosome separation (Mayer et al., 1999). *Sciara* male meiosis I exhibits one of the few known natural occurrences of a monopolar spindle (Metz, 1933 and 1936), that has also been reported for the beetle *Micromalthus* (Scott, 1936) and in the gall midge *Mycophila* (Fux, 1976; Camenzind and Fux, 1977).

Similar to *Sciara* male meiosis I, we observed that spermatocytes in meiosis II first form a single half-spindle from the single centrosome. However, later in metaphase a second yet shorter anastral half-spindle forms on the other side of the metaphase plate, as deduced by Fuge (1999) in fixed spermatocytes from the sciarid *Trichosia*. The two poles of the asymmetric spindle in *Sciara* meiosis II differ, as γ-tubulin, centrin and other centrosomal proteins are only at the single centrosome but not at the anastral pole (Esteban et al., 1997). In addition to the *Sciara* secondary spermatocytes studied here, monastral bipolar spindles have been documented in other systems. They can be induced in crane flies (Dietz, 1966; Bastmeyer et al., 1986; Steffen et al., 1986) or by mutation in *Drosophila* (Sunkel and Glover, 1988; Llamazares et al., 1991; Wilson et al., 1997), and they can occur normally, as in *Drosophila* oocytes (Theukauf and Hawley, 1992; McKim and Hawley, 1995; Endow and Komma, 1998) and in parthogenetic *Sciara* embryos (de Saint Phalle and Sullivan, 1998).

How does the anastral half-spindle form in *Sciara* meiosis II? In the absence of centrosomes, other pathways exist to nucleate spindle microtubules. For example, bipolar spindles form in the asterless *Drosophila* mutant (Bonaccorsi et al., 1998). Acentrosomal microtubules assembled through a chromatin-dependent pathway are necessary and sufficient for spindle formation in all cell types (Karsenti et al., 1984; Heald et al., 1996; Khodjakov et al., 2000). Microtubules can nucleate around DNA and form bipolar spindles by microtubules motors (Heald et al., 1996 and 1997; Merdes and Cleveland, 1997; Hyman and Karsenti, 1998). Moreover, chromosome-bound RCC1 creates a Ran-GTP gradient near the chromosomes to promote microtubule nucleation and stabilization by spindle assembly factors (Cavazza and Vernos, 2016). In addition, kinetochores can serve as secondary microtubules organizing centers (McGill and Brinkley, 1975; Mitchison and Kirschner, 1984). Microtubules initiated by the chromosomes are stabilized when their plus ends bind to kinetochores while their minus ends are bundled by dynein into a bipolar spindle (Khodjakov et al., 2000; Karsenti and Vernos, 2001). Our data suggest that these additional pathways to form spindle microtubules are less efficient and take longer than use of the centrosome as a microtubules organizing center that initially creates a monopolar half-spindle at the beginning of meiosis II in *Sciara* spermatocytes.

Why is the asymmetric monastral bipolar spindle found in meiosis II but not in meiosis I? Perhaps the checkpoint protein Mad2 found at the X dyad lengthens metaphase sufficiently to allow the second half-spindle to form. In contrast, chromosomes in meiosis I do not have Mad2 localized to their centromeres. Moreover, spermatocytes in meiosis I lack metaphase, which is when the spindle assembly checkpoint would normally act. Another possibility is that the smaller volume of the nucleus in meiosis II than in meiosis I (Table S1) could facilitate the bundling and focusing of the acentrosomal, chromatin-dependent microtubules into the non-centrosomal half spindle. A corollary to this idea is that the paternal chromosomes on the monopolar spindle of meiosis I might have inactivated the chromatin-dependent microtubule polymerization pathway, which would remain active on the maternal chromosomes in meiosis II. Furthermore, might this be related to the meiotic drive gene segregation distorter in *Drosophila* that has a RanGAP duplication (Larracuente and Presgraves, 2012)?

### Chromosome Movement on Meiotic Spindles in *Sciara* Spermatocytes

In male *Sciara* meiosis I how the maternal chromosomes move to the single centrosome and how the paternal; chromosomes move away from the single centrosome has remained a mystery. Previously, c-banding revealed that the centromeres on the paternal chromosomes orient towards the single centrosome, though their apparent movement is away from the single centrosome (Abbott et al, 1981). If the paternal chromosomes move away from the single centrosome, one would expect the non-centrosomal chromosome ends to follow rather than lead the direction of motion due to viscous retardation by the cytoplasm. However, this is not the case. Moreover, breaking a chromosome arm by x-irradiation suggested that there is no neocentromere nor unique attachment of the non-centrosomal end of the chromosome to pull or anchor it to structures in the cell distal to the single centrosome (Abbott et al., 1981). It is possible that polar ejection forces (Rieder et al, 1986) created by chromokinesins on the arms on the arms account for “backward” orientation of the paternal chromosomes in male meiosis I where the centromere lags behind the apparent direction of movement away from the single monopole.

Electron microscopy revealed kinetochores on the paternal but not on the maternal chromosomes in meiosis I of *Sciara* spermatocytes (Kubai, 1982). Conversely, electron microscopy detected kinetochores on the maternal but not the paternal chromosomes in meiosis I of spermatocytes from the sciarid *Trichosia* (Fuge, 1994 and 1997), consistent with the lack of paternal chromosome orientation seen in *Trichosia* primary spermatocytes by light microscopy (Amabis et al., 1979). It is possible that which chromosome sets have kinetochores differs between *Sciara* and *Trichosia*. Alternatively, these disparate observations may be reconciled if there were differential timing for kinetochore occurrence in the two sets of chromosomes. For example, kinetochores might form first on the maternal chromosomes, allowing them to be pulled to the single pole at which point their kinetochores would degenerate; subsequently, kinetochores might form on the paternal chromosomes that back away from the single pole. This is reminiscent of an earlier hypothesis that the maternal chromosomes move before the paternal chromosomes in *Sciara* male meiosis I (Kubai, 1987).

Kinetochores would orient chromosomes of each set with their centromeres closest to the single pole, but do the kinetochores play a role for chromosome movement on the monopolar spindle of meiosis I? A “pacman-like” action of the kinetochore to shorten the spindle microtubules at their kinetochore attachment point is plausible for the maternal set of chromosomes, but not for the paternal set where they back away from the single pole. Might the microtubules on the paternal chromosomes be lengthening rather than shortening or might other forces such as “polar winds” (Rieder et al, 1986) push the paternal chromosomes away from the single pole? In our observations reported here we never saw significant chromosome movement in meiosis I in living spermatocytes, raising the possibility that the chromosomes might not move on the monopolar spindle.

In meiosis II of *Sciara* spermatocytes, Metz suggested that the X dyad had a separate time-clock and moved precociously from the metaphase plate to the pole (Metz, 1938). Our observations reported here with polarization optics of living spermatocytes demonstrate that the conclusion by Metz is not correct. We never saw the X dyad on the metaphase plate in meiosis II; conclusions similar to ours have been deduced by observations on fixed spermatocytes of *Trichosia* (Fuge, 1999). We saw that the X dyad remains near the centrosome after meiosis I and moves outward with it as the spindle forms in meiosis II. Moreover, amorphous material between the X dyad and the centrosome was visualized in meiosis II spermatocytes by electron microscopy, perhaps serving as a glue to tether the X dyad to the single centrosome. Although the distance increases between the X dyad and the metaphase plate during metaphase, it may reflect pulling on the X by the juxtaposed centrosome rather than active movement of the X dyad on the spindle. Moreover, a well-formed kinetochore was not seen by electron microscopy on the X dyad, in contrast to the chromosomes on the metaphase plate. Although a kinetochore was not visible on the X dyad, the checkpoint protein Mad2 associated with the X dyad centromeres.

### Chromosome Identification by *Sciara* Spermatocytes

How does the cell identify the paternal chromosomes for elimination in meiosis I and the X dyad for non-disjunction in meiosis II? Our data show that the paternal chromosomes contain Mad2 throughout their length in the anaphase-like configuration on the monopolar spindle in meiosis I of *Sciara* spermatocytes. Normally the checkpoint protein Mad2 is localized to kinetochores in metaphase of mitotic cells until they attach to spindle microtubules (Waters et al., 1999; Logarinho et al., 2004). In male meiosis of other systems, Mad2 remained at most kinetochores until metaphase II (Kallio, 2000). The monopolar spindle of meiosis I in *Sciara* spermatocytes lacks metaphase when Mad2 would normally associate with kinetochores. Instead, the location of Mad2 decorating the entire length of the paternal chromosomes suggests it might have a second role and help to identify the imprinted paternal chromosomes. There is precedent for centromeric proteins having a second role, such as the chromosomal passenger complex that is shed from centromeres to the spindle midzone where it functions in cytokinesis (Carmena et al, 2012). Besides Mad2, some other differences between the maternal and paternal chromosomes in *Sciara* male meiosis I. have also been observed. There is acetylation of histones H3 and H4 only on the maternal chromosomes (Goday and Ruiz, 2002), and only the eliminated paternal chromosomes show methylation of histone H3 at lysine 4 (Greciano and Goday, 2006). It is unknown if these are the cause or consequence of chromosome imprinting.

In meiosis II of *Sciara* spermatocytes the X dyad destined for non-disjunction is marked by the “controlling element” (Crouse, 1960). This locus can be moved by translocation to an autosome that then exhibits non-disjunction in male meiosis II, whereas the translocation X lacking this locus now segregates to the two poles of the bipolar spindle (Crouse, 1960). Therefore, the controlling element acts in *cis* on the chromosome where it resides, marking it for non-disjunction in meiosis II of *Sciara* spermatocytes. Because the X dyad lacks a well-defined kinetochore, it is tempting to speculate that the controlling element may prevent kinetochore formation or play a role in production of the amorphous substance between the X centromere and the single centrosome. The same controlling element eliminates the “extra” X chromosome(s) from the embryonic soma by an apparently different mechanism where the X moves to the metaphase plate but the sister chromatids fail to separate (de Saint Phalle and Sullivan, 1996).

With the assembly of the *Sciara* genome (Urban et al 2021), future studies may elucidate the molecular details of the imprint that distinguishes the paternal chromosomes for elimination in meiosis I and the mechanism of action of the controlling element for X dyad non-disjunction in meiosis II of *Sciara* spermatocytes.

## MATERIALS AND METHODS

### Fly Stocks

The *Sciara* Stock Center at Brown University provided Sciara coprophila (syn. Bradysia coprophila Lintner) strains 7298 and 6980, which both carry the dominant marker Wavy on the X’ chromosome; stock 6980 also carries the recessive swollen vein marker on the X chromosome.

### Live Observations

In *Sciara* the male meiotic divisions occur during pupation when the omatidia of the eye become pigmented. Pupae were selected when the fraction of black omatidia was between 1/4 and 2/3.

For DIC observations, pupae were washed in distilled water, briefly in ethanol and lightly dried on filter paper. Super-clean slides and coverslips were used which had been sonicated for 30 minutes in Dynasoap solution, rinsed with tap and distilled water and ethanol. They were then stored in ethanol which was removed immediately before use by flaming with an alcohol lamp and then air cooled. The spermatocytes are very sensitive to temperature; therefore, the dissections and microscopic observations were done in a room at 19^0^C with high humidity. Each pupa was dissected under lightweight Nujol oil, taking care to not rupture the gut or Malpighian tubules which would liberate toxic substances. Each of the two testes was moved to an area of oil without material from the rest of the carcass and then picked up with forceps underneath the testis for transfer to a drop of Nujol oil on a super clean coverslip where it was dragged with the forceps to spread out the cells. A super-clean slide was placed on the coverslip, separated with a spacer of pieces of broken coverslip at the corners of the coverslip containing the preparation and was tacked down to the slide with molten VALAP (1:1:1 vaseline, lanolin, paraffin) at the corners. Visibility of the preparation improved after 10 minutes when the cells had flattened onto the coverslip. Three glass plates were used as a heat filter for observations with DIC and polarization optics.

For LC-PolScope observations, pupae were dissected in tricine insect buffer (70 mM NaCl, 45 mM sucrose, 25 mM Tricine (N-Tris (hydroxymethyl) - methyl glycine, pH 7.2), 15 mM NaHCO_3_, 8 mM KCl, 8 mM MgCl_2_.6H_2_0; Begg and Ellis, 1979) using a dribble ball action with the forceps to remove fat from the testes. The isolated testes were ruptured under Voltalef 10s oil (Ugine Kuhlmann, Paris, France) in a ring of Vaseline on a super-clean glass coverslip with a drop of molten VALAB (1:1:1 vaseline, lanolin, and bees wax) or VALAP at each corner. The coverslip was placed on a glass slide between spacers of coverslip slivers. A second microscope slide was then pressed down on the spacers compressing the VALAB, which attached the coverslip to the second slide. Spermatocytes survived in these preparations for up to 2.5 hours as shown by a sequence of images from meiosis II metaphase through telophase. However, except for the transition from metaphase II into telophase II the cells were very light sensitive and seldom continued meiotic progression once they were imaged with the LC-PolScope despite the use of a reflective heat filter (HotMirror). This limited capturing long sequences to the progression from metaphase II to telophase II and into budding. The light was less intense with standard polarization and DIC optics where sequences from prophase II through metaphase II were also observed in the living cells.

Two variations on the preparation described above were used to image the specimen in both polarized light and epifluorescence. The DNA was stained with Hoechst 33342 (Molecular Probes; cat. #H-1399; Invitrogen, Carlsbad, CA) by adding Hoechst to the tricine insect buffer used for dissection at a final concentration of 1 μg/ml for 5 minutes. The testis was washed in plain buffer before being ruptured under oil. The other variation was the “Oil prep” where the pupa was dissected directly under Voltalef oil to eliminate any effect from the insect buffer. The results were the same with both methods and did not improve capturing long sequences through meiosis.

Polarized light images were obtained with the LC-PolScope (Cambridge Research and Instrumentation, Woburn, MA; Oldenbourg and Mei, 1995) as described earlier (LaFountain and Oldenbourg, 2004; Janicke et al., 2007). The LC-PolScope is a polarized light microscope equipped with a computer-controlled liquid crystal universal compensator. Specimens were imaged with a 60X/1.4 NA plan apochromat oil immersion objective and achromat oil immersion condenser at five predetermined polarization states. The images were recorded with a CCD camera (Retiga EXi FAST, Q-Imaging) and stored as TIFF files. Using the 5 raw LC PolScope images, an orientation independent retardance image representing the birefringence of the specimen was obtained (Shribak and Oldenbourg, 2003). Testes stained with Hoechst were also imaged (with the same lens and CCD camera) by epifluorescence using a Nikon UV-1A filter cube.

### Calculations of Nuclear Volume

To get a numerical estimate, the cell area and nucleus area (area delineated by the Hoechst-stained chromosomes) were measured with ImageJ (Rasband, W.S., ImageJ, U. S. National Institutes of Health, Bethesda, MD, http://rsb.info.nih.gov/ij/, 1997-2006) in a cross section and the ratio was computed individually for each of 11 cells. The mean of these computed proportions was 0.59 (59%) of the cell area (Supplemental Table S1). The proportion of the cell volume occupied by the nucleus was roughly estimated by raising the proportion of the area occupied by the nucleus to the 3/2 power. The resulting estimate of the volume occupied by the chromosomes is 0.45 or 45% of the volume of the cell. The cell cross-section area was the same in PI and PII (p = 0.99)(F Test for equal variance: F = 0.9844 alpha= 0.01, t-Test for equal means at alpha = 0.01 t = -0.9987 t critical (2 tailed) = 2.8453). The mean of the proportions of nucleus area to cell area for PII was 0.21 or 21% (Supplemental Table S1). The proportion of the cell volume occupied by the nucleus estimated as above for PI was 0.09 or 9% of the volume of the cell.

### Immunofluorescence

Pupae were dissected in tricine insect buffer, disrupted in a small amount of this buffer on a slide treated with CellTak (Biopolymers Farmington, CT), Polylysine or APTES (3-aminopropyltriethoxysilane)(Sigma, St. Louis, Mo.) for adhesion. For Mad2 staining, cells were fixed by immersion in methanol at -20^0^C for 10 minutes followed by fixation in 4% EM grade paraformaldehyde (Electron Microscopy Sciences, Hatfield, PA) for 10 minutes followed by permeabilization and blocking in PBTA Blocking Solution (PBS 1X, BSA 3%, Triton-X100 0.3%, sodium azide 0.03%) for one hour. The R793 antibody against *Drosophila* Mad2, kindly provided by Tom Maresca from Gregory Rogers (University of North Carolina - Chapel Hill), was used for staining at 1:100 in PBTA (PBS 1X, BSA 1%, Triton-X100 0.1%, sodium azide 0.03%) and mounted in Vectashield with propidium iodide (Vector Laboratories, Burlingame, CA) to stain the DNA (double stain), then mounted in Vectashield (Vector Laboratories, Burlingame, CA) without propidium iodide. When co-staining for tubulin with a monoclonal antibody, instead of propidium iodide, Hoechst dye was used at a concentration of 1 μg/ml. Immunofluorescence images were obtained using a Zeiss LSM confocal, 63X Zeiss lens, a Chameleon laser for 2 photon, and HeNe and Argon 488 lasers. Collimation of Chameleon and other lasers was not complete using microscope adjustments, and required 1 um offset of blue (2 photon) stack.

### Electron Microscopy

Testis preparation and electron microscopy was done as described earlier (Kubai, 1982 and 1987), using a Zeiss 10A electron microscope at 80 kV with 70 mm film (No. 613, Chemco Photoproduction, Glen Cove, NY).

### Western Blots

Western blots against Mad2 detect a specific band at 25 kD in *Drosophila* (Buffin et al., 2007), and the R793 antibody detected the same size band in *Sciara* extracts as in *Drosophila* extracts. For Western blots, 50 μl packed *Drosophila* or *Sciara* embryos or 6 larvae were mashed with a pestle in an Eppendorf tube containing 30 μl distilled water, 60 μl 10% SDS, 90 μl 50% glycerol and 30 μl DTT. Following a standard protocol (Sambrook et al., 1989), 150 μl 2X loading buffer was added, and the sample was boiled for 5 minutes and spun down. The supernatant was loaded on a 10% SDS-polyacrylamide gel with a 4% stacking gel and run at 188V for 45 minutes. It was transferred to nitrocellulose paper at 4^0^C for 45 minutes with 0.35 amp constant current and blotted 30-60 minutes in milk/TBST. The sample was incubated with diluted R793 anti-Mad2 antibody at 4^0^C overnight or 1 hour at room temperature, washed 3 times 6 minutes each in milk/TBST, and incubated with the secondary antibody (1:300 dilution of donkey anti-rabbit) 1 hour 45 minutes at room temperature followed by 3 washes of 6 minutes each in milk/TBST. The membrane was treated with equal volumes of solution A and B of the Perkin Elmer ECL kit (Waltham MA) for 1 minute at room temperature.

## ACKNOWLEDGEMENTS

Thanks to Tom Maresca for providing the Mad2 antibody from Greg Rogers, to Alison Singer, Janell Johnson and Stephen Doris for performing the Western blots, and to R. Bruce Nicklas and James R. LaFountain Jr for technical advice on *Sciara* spermatocyte preparations This research was supported by NIH EB 002583 (to RO), NIH GM 35929 and NIH GM121455 (to SAG), and NIH fellowship GM08514 to SAG that funded her sabbatical. Thanks to Sharyn Endow and R. Bruce Nicklas for the hospitality of their labs at Duke University during this sabbatical.

## DEDICATION

This paper is dedicated in memory of our mentor and colleague Shinya Inoué (1921-2019), a pioneer in polarization microscopy, whose work to advance this methodology laid the foundation for the research described here.

---

**Suppl. Figure S1.**
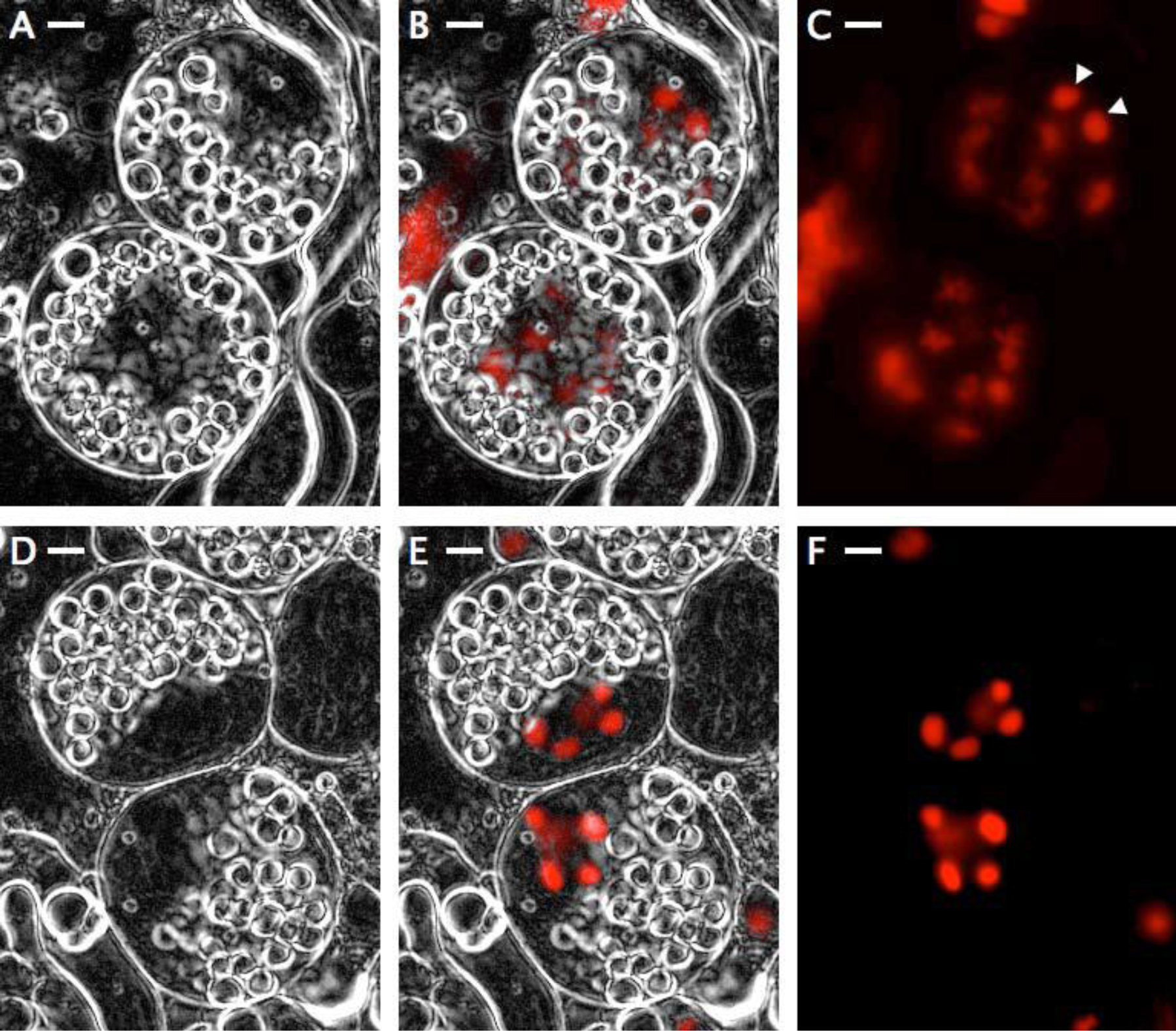
Prophase I and II (PI and PII). Live LC-PolScope images and epifluorescence images of DNA stained with Hoechst. In PI, the chromosomes are loosely condensed except the L chromosomes, which are more tightly condensed (arrowheads in C). In PII the chromosomes are more tightly condensed than in PI and the L chromosomes are no longer differentially condensed; also, the PII nucleus occupies a smaller volume than PI (see Supplemental Table 1). All bars = 2 μm. **A-C:** 2 PI cells. **A.** LC-PolScope image. **B.** LC-PolScope image combined with Hoechst image of DNA (red). **C.** Image of DNA stained with Hoechst. Arrows mark the L chromosomes in the upper nucleus where they are in the plane of focus **D-F:** 2 PII cells. **D.** LC-PolScope image. **E.** LC-PolScope image combined with Hoechst image of DNA (red). **F.** Image of DNA stained with Hoechst.

**Supplemental Table 1.** Size comparison of prophase I and prophase II

|  | **Cell area** | **Nucleus area** | **Ratio nucleus/cell** |
| --- | --- | --- | --- |
| **PI** | 18871+/-1314 | 10920 +/-594 | 0.5912 +/-0.0279 |
| **PII** | 20730 +/-1324 | 4329 +/-500 | 0.2060 +/-0.0157 |
| Difference |  |  | 0.3852 +/-0.032 |

All data are means +/-STE, n=11 in pixels calculated with ImageJ Cell areas to the nearest pixel, proportions to 4 significant figures Statistics computed with EXCEL Data Analysis Cell area PI = Cell area PII (F test indicates equal variance $\alpha=0.01$ , t-Test (2 tail equal variance) indicates equal means $\alpha=0.01$ ) Ratio nucleus/cell PI & PII unequal (F test indicates unequal variance $\alpha=0.05$ , t-Test (2 tail unequal variance) indicates unequal means $\alpha=0.001$ )

